# Design of Cytotoxic T Cell Epitopes by Machine Learning of Human Degrons

**DOI:** 10.1101/2023.08.22.554289

**Authors:** Nicholas L. Truex, Somesh Mohapatra, Mariane Melo, Jacob Rodriguez, Na Li, Wuhbet Abraham, Deborah Sementa, Faycal Touti, Derin B. Keskin, Catherine J. Wu, Darrell J. Irvine, Rafael Gómez-Bombarelli, Bradley L. Pentelute

## Abstract

Antigen processing is critical for producing epitope peptides that are presented by HLA molecules for T cell recognition. Therapeutic vaccines aim to harness these epitopes for priming cytotoxic T cell responses against cancer and pathogens, but insufficient processing often reduces vaccine efficacy through limiting the quantity of epitopes released. Here, we set out to improve antigen processing by harnessing protein degradation signals called degrons from the ubiquitin-proteasome system. We used machine learning to generate a computational model that ascribes a proteasomal degradation score between 0 and 100. Epitope peptides with varying degron activities were synthesized and translocated into cells using nontoxic anthrax proteins: protective antigen (PA) and the N-terminus of lethal factor (LF_N_). Immunogenicity studies revealed epitope sequences with a low score (<25) show pronounced T-cell activation but epitope sequences with a higher score (>75) provide limited activation. This work sheds light on the sequence–activity relationships between proteasomal degradation and epitope immunogenicity, through conserving the epitope region but varying the flanking sequence. We anticipate that future efforts to incorporate proteasomal degradation signals into vaccine designs will lead to enhanced cytotoxic T cell priming by vaccine therapeutics in clinical settings.

## Main Text

Vaccines are transforming human health through equipping a patient’s own immune system to defend against cancer and pathogenic disease^1,2^. The components of a vaccine aim to mimic the biological processes associated with acquiring natural immunity, by generating immune cell populations that recognize tumor- and pathogen-specific epitopes^3^. Vaccine formulations that prime new and existing T cell populations offer the promise of not only eliminating affected cells, but also for providing long-term protection through immune-memory responses^4^. Despite this promise, the design and development of vaccine epitopes that provide robust priming of cytotoxic T cells has thus far proved difficult. A key challenge is thought to be due to insufficient epitope processing and presentation, which leads to reduced vaccine efficacy and has thus far precluded FDA approvals for T cell vaccines^5^.

Developing a vaccine is a multistep process that includes characterizing molecular sequences from tumor, bacterial, or viral proteins, selecting one or more immunogenic epitopes, and administering the epitopes within a longer polypeptide sequence as a peptide, RNA, DNA, or viral vector sequence (Fig. 1a)^6–10^. Epitopes aimed at priming cytotoxic T cell responses are shorter peptides (8–9 amino acids) that load into class I HLA molecules, followed by transport to the cell surface for antigen presentation^11^. Although computational tools have enabled the prediction of high-affinity HLA-binding epitopes that favor loading^12–14^, intracellular processing of the longer polypeptides is essential for producing these epitopes^15–18^. To date, the rules for predicting the hydrolytic propensity of a given epitope have remained elusive^19,20^.

**Fig. 1.**
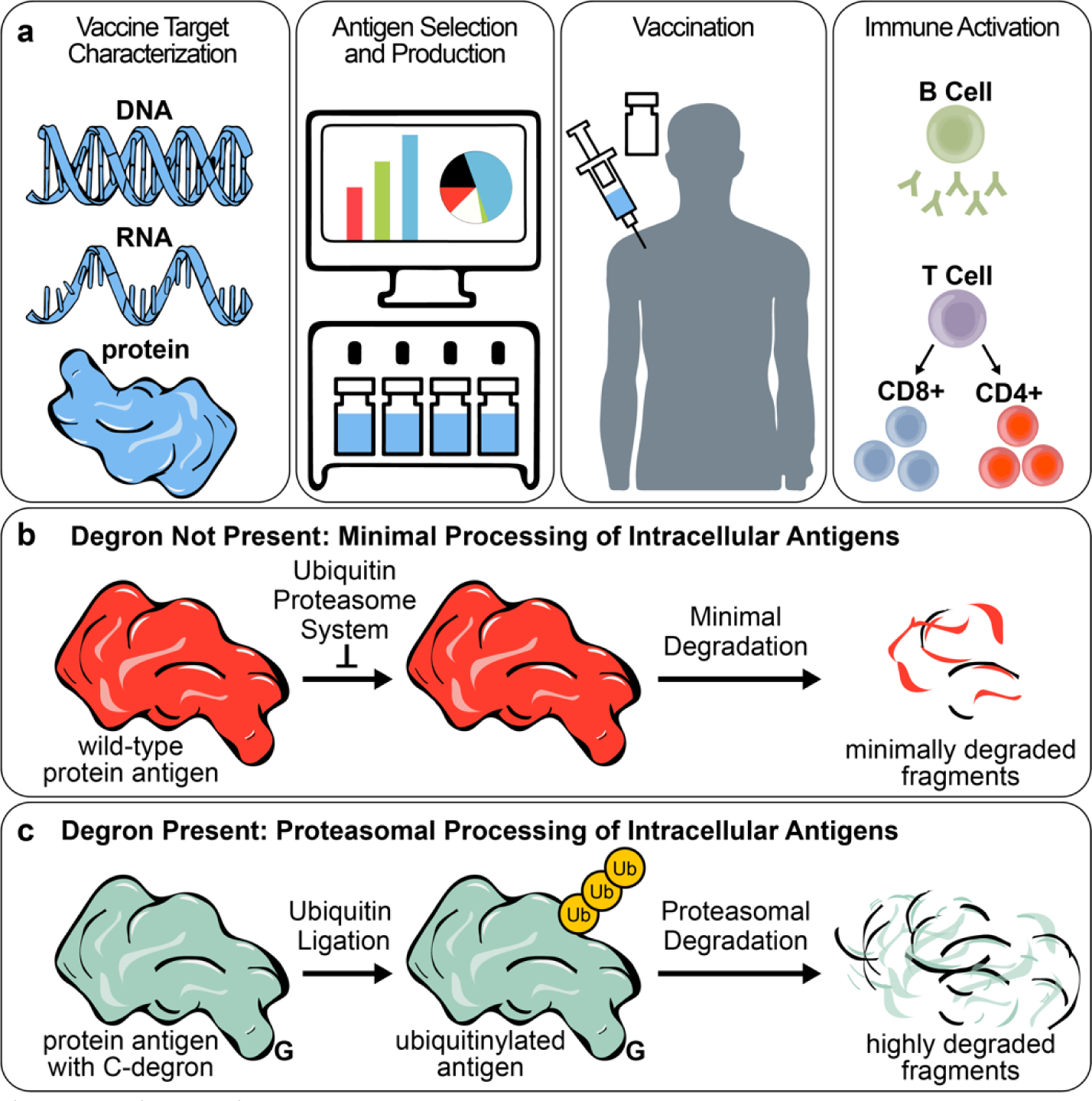
Vaccine Design with Human Degrons. **a**, Schematic for vaccine design, which is enabled by target characterization through DNA, RNA, or protein sequencing; selection and production of cytotoxic T cell epitopes; administration of the vaccine components; and immune priming of epitope-specific T and B cells. **b**, Epitopes without a degron sequence undergo minimal processing. **c**, Epitopes containing C-terminal degron (C-degron) sequences undergo ubiquitin-mediated proteasomal degradation, in which the resulting fragments enable antigen presentation by HLA molecules and the priming of cytotoxic T cells.

Intracellular degradation of peptides and proteins is prevalent in cellular metabolism, and may provide rules for designing vaccine epitopes and other therapeutic polypeptides^21^. The degradation is based on short peptide sequences, called degrons, that signal the proteasome for protein degradation^22^. In 1986, Varshavsky and co-workers discovered that a single amino acid at the N-terminus can dictate the propensity of proteasomal degradation, which is now known as the “N-end rule”^23^. In 2015, we similarly observed that a single D-amino acid can abrogate proteasomal degradation, perhaps due to the absence of ubiquitin ligases that can recognize D-amino acids^24^. Moreover, Elledge and co-workers demonstrated longer stretches of amino acids, up to 23 residues in length, adopt sequence patterns at either the N or C terminus that greatly influence the magnitude of proteasome degradation^25–30^. These protein degradation studies show the influence of degron sequences on regulating proteasomal activity, but also suggest that the absence or presence of a degron may even influence the immunogenicity for a given vaccine epitope (Fig. 1b,c)^31^.

In this work, we set out to uncover vaccine design rules that can infer epitope immunogenicity from proteasomal degradation activity. We used machine learning to create a degron prediction model, in which the model enables interpretation of complex sequence patterns and their proteasomal stabilities. The training data comprises the proteasomal stabilities of 22,564 sequences from the C-termini of human proteins, and reflects the propensity of protein degradation across the human proteome^29^. The resulting model ascribes a relative degradation score between 0 and 100, which is determined from the C-terminal residues of a peptide or protein sequence. The model does not include stability data from N-terminal sequences because multiple non-proteasome mechanisms are known to hydrolyze the N-terminus, including N-terminal trimming from an endoplasmic reticulum aminopeptidase (ERAP) 1 or 2^31,32^.

To validate our C-degron model, we used a protein delivery system that efficiently transports epitope peptides into antigen presenting cells to ensure cytosolic delivery and provide access to the proteasomal degradation machinery. We used two nontoxic anthrax proteins for epitope translocation: protective antigen (PA) and the N-terminus of lethal factor (LF_N_). Incorporating C-degron peptides at the C-terminus of LF_N_ enabled validation of predicted proteasomal degradation activity by Western blot analysis and a series of T cell proliferation assays. These studies show that combining C-degron sequences with epitope peptides favors proteasomal degradation and, in turn, maximizes epitope immunogenicity.

## Results

### Machine learning for prediction of human degrons

Previously, we demonstrated that deep neural network models can be trained to relate peptide sequence to biological activity, and to support design loops for developing novel bioactive peptide sequences^33^. We train 1-dimensional convolutional networks on peptide sequences, by representing the monomer identity as a fingerprint that reflects the chemical structure (Fig. 2a,b). Here, we used these methods to associate amino acid sequences with their degradation propensity.

**Fig. 2.**
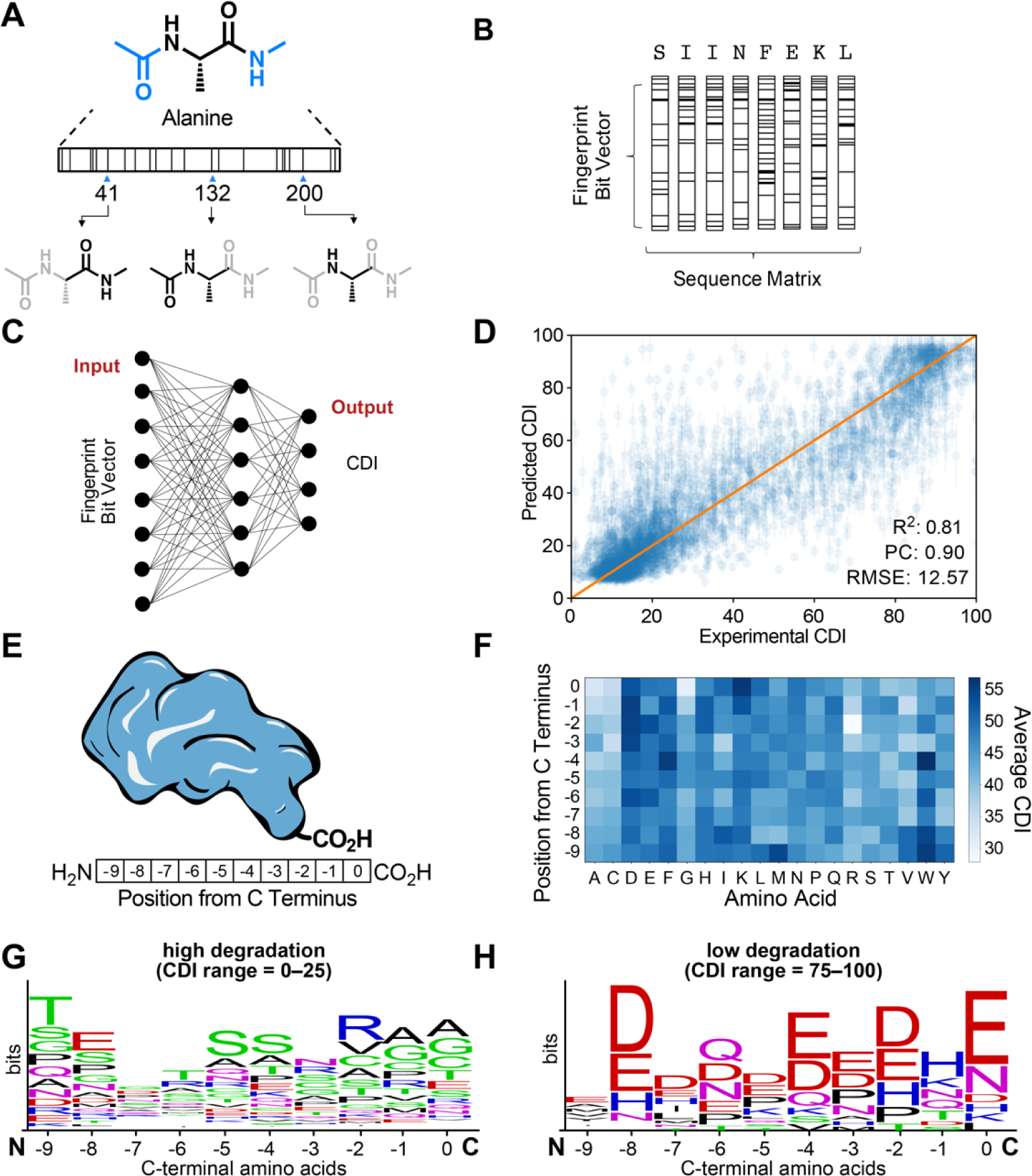
Machine Learning of Human Degron Sequences. **a**, Fingerprint representation of a single amino acid. **b**, Sequence matrix representation for the SIINFEKL epitope of Ovalbumin_257-264_. **c**, Model architecture with sequence matrix inputs and outputs for predicting C-terminal Degradation Index (CDI). **d**, Observed versus predicted parity plots for linear regression analysis of CDI. Training data were split into three subsets for model training (60%), cross-validation (20%), and testing (20%). Inset text and graphics indicate key performance metrics: R^2^, Pearson’s correlation (PC), root-mean-squared error (RMSE), and solid orange line, y = x line. (E–H) Analysis of randomized sequences (30,000) based on the C-terminal amino acids on a protein. **e**, Illustration of the C-terminal residues (10 amino acids) that influence proteasomal stability. **f**, Heat map plot showing the C-terminal amino acids (10 amino acids) and average CDI values (ranging from 30 to 55). **g–h**, Sequence logo plots of recurrent amino acids at corresponding positions, which were plotted across CDI values for lower (CDI = 0–25) and upper (CDI = 75–100) quartiles.

The training data were obtained from stability index studies previously described by Elledge and coworkers, comprising a plasmid library of DNA-encoded peptides from the human proteome^29^. The peptides include the 23 residues from the C-termini of human protein sequences which are fused to the green fluorescent protein (GFP). In the prior work, transfection of these DNA libraries into mammalian cells demonstrated protein degradation from the GFP expression levels. The experimental data revealed a distribution of fluorescent cells among numeric ‘bins’ (e.g., bin1, bin2, bin3, and bin4). The fraction of cell populations in the lower bins (e.g., bin1) indicate reduced fluorescence intensity associated with GFP degradation; the fraction of cell populations in the higher bins (e.g., bin4) indicate high fluorescence intensity associated with intact GFP.

We developed a linear equation that condenses the proteasomal stability data to a single score, which we call the C-terminal degron Index (CDI). This score is calculated from a linear combination of two parameters: bin population data (e.g., bin1, bin2, bin3, and bin4), which was obtained from Elledge and coworkers^29^; and exponential coefficients (e.g., 0, 1, 10, and 100), which reflect the exponential scale of the original data (i.e., flow cytometry). The resulting score relates degradation propensity to a numerical CDI that ranges from 0 to 100. Interpreting the CDI for a given sequence is straightforward: a CDI value in the lower quartile (i.e., 0–25) reflects pronounced degradation; a CDI value in the upper quartile (i.e., 75–100) reflects limited degradation. The CDI also enables comparisons of closely related values (e.g., 20 vs 25), and their corresponding degradation propensity.

To prepare the input data for machine learning, we calculated CDI values across the human proteome using the sequence data from Elledge and co-workers (22,564 sequences). The machine learning model was established using 100% of the input sequences and CDI values; however, the model was trained, validated, and tested using the following three randomized subsets of the data: 60%, 20%, and 20% (Fig. 2c). On a 20% subset, we calculated several measures of statistical significance by fitting the data to the CDI model. The resulting analysis showed: root-mean-squared error (RMSE), CDI_RMSE_ = 12.97 ± 0.11; linear regression R-squared value, R^2^ = 0.796 ± 0.003; and Pearson correlation efficient, ρ = 0.896 ± 0.002 (Fig. 2d). These values show that the CDI model can reasonably ascribe degradation propensity to a given sequence located at the C-terminus of a peptide or protein.

We interrogated the model by generating a randomized sequence library and analyzing sequence trends based on CDI. The library was obtained from 30,000 randomly-generated sequences that vary at the C terminus (i.e., 10 residues) but maintain a constant N terminus (Fig. 2e). We generated a heat map to evaluate sequence trends that emerge for the ten C-terminal residues (i.e., positions 0 to -9) (Fig. 2f). We also generated sequence logo plots for two key populations, in which one plot reflects the lower quartile (CDI: 0–25, Fig. 2g) and the other reflects the higher quartile (CDI: 75–100, Fig. 2h).

The heat map and sequence logo plots reveal identities and positions of amino acids that favor degradation (Fig. 2f,g). Consistent with prior experiments, the model predicts that a Gly residue located at position 0 or -1 contributes substantially to degradation propensity (i.e., CDI ≤ 30); the model also predicts that a Gly residue located further from the C terminus (i.e., position -6 or -7) also favors proteasomal degradation but to a lesser extent^27^. The plots show that other amino acid residues are also predicted to favor degradation (i.e., CDI ≤ 40): an Ala residue at position 0, -1, -2, -4, -5, -6, or -7; a Cys residue at position 0, -1, -2, -3, -4, -5, or -6; an Ile residue at -3 or -6; an Arg residue at position 0, -1, -2, -3, -6, -8, or -9; a Ser residue at position - 3 or -8; a Thr residue at position 0, -4, or -5; and a Val residue at position 0, -1, -2, -3, -5, -6, or - 7. These amino acid residues are thought to reflect substrates of ubiquitin ligase enzymes, which mediate ubiquitin ligation and subsequent proteasomal degradation.

The model also predicts amino acid residues that mostly do not favor degradation (i.e., CDI > 40); these residues include Asp, Glu, Asn, Gln, Phe, Pro, His, Lys, Leu, Met, Trp, and Tyr. The heat map and sequence logo plots shed light on the identities and positions of residues that do not favor degradation (Fig. 2f,h). Nonetheless, CDI is determined from an overall sequence rather than the individual residues and, therefore, a sequence may still give a low CDI while containing one or more residues that do not favor degradation.

### Predicted degrons regulate proteasomal degradation

Bacterial toxin proteins have previously been shown to enable protein degradation studies, because these studies are otherwise notoriously challenging without ensuring cytosolic delivery^24,34^. Here, two anthrax proteins were used for cytosolic epitope delivery: PA and LF_N_. Previously, PA/LF_N_ have been shown to enable cytosolic delivery of cytotoxic T cell epitopes^35,36^. The proteins mediate protein translocation through PA binding to a transmembrane protein receptor, either tumor endothelial cell marker (TEM8) or capillary morphogenesis protein 2 (CMG2), and mediating the delivery of lethal and edema factors into the cytosol of mammalian cells^37,38^.

In the current study, peptides were conjugated to the C-terminus of LF_N_ using sortase-mediated ligation (Fig. 3a)^39^. These LF_N_-peptide conjugates were co-administered with the pore-forming protein, PA, enabling cytosolic delivery through a multistep mechanism (Fig. 3b)^40^. The PA-mediated translocation mechanism is well established, which includes: (1) binding to receptors on mammalian cells; (2) undergoing cleavage to give 20-kDa and 63-kDa fragments, called PA_20_ and PA_63_, followed by PA_63_ assembling to form annular heptamers (PA_63_)_7_; (3) further assembling with LF_N_ molecules; (4) entering the cell endosome; and (5) rearranging for insertion into the endosomal membrane for LF_N_ translocation into the cell.

**Fig. 3.**
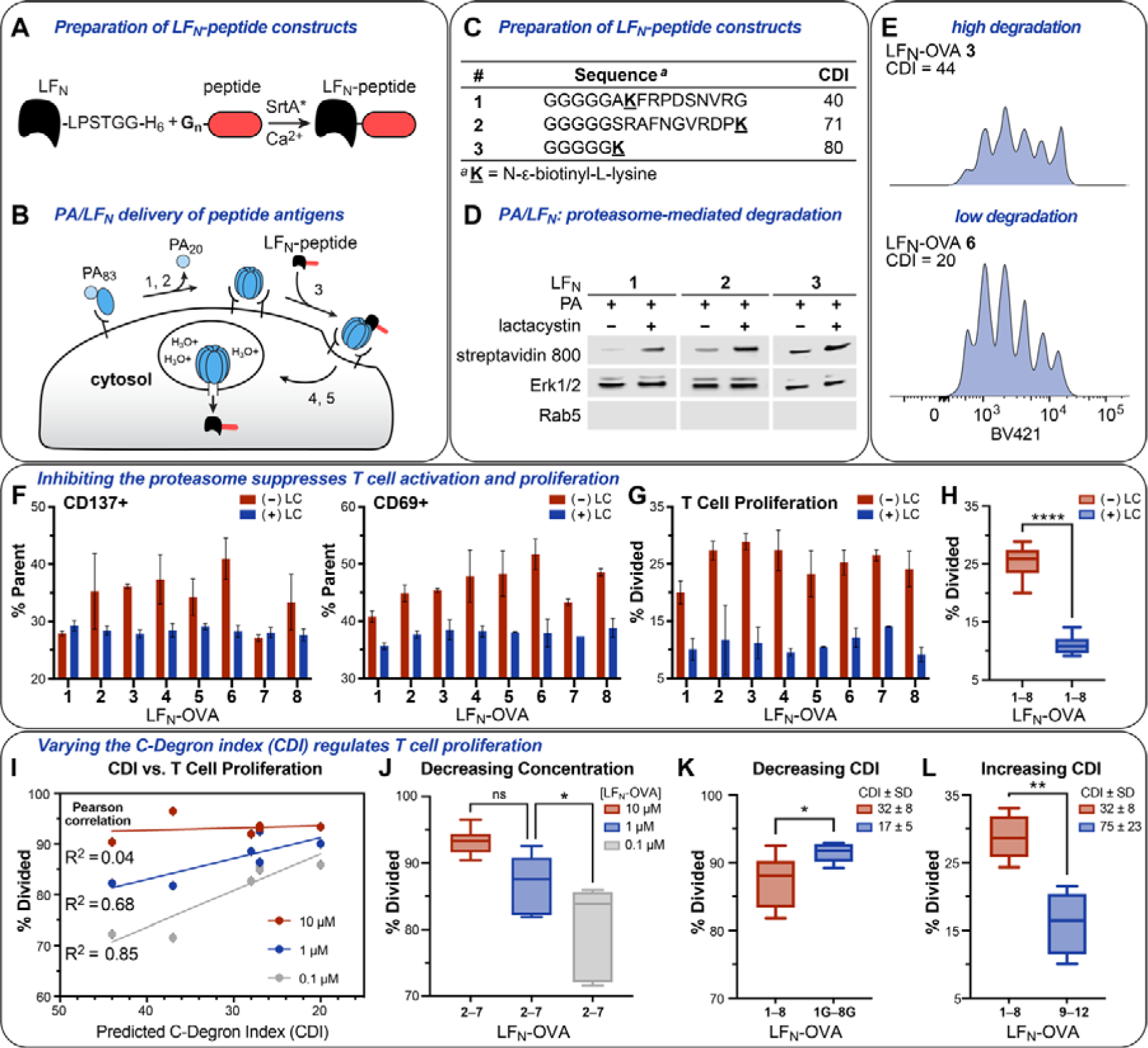
Proteasomal stability reflects the magnitude of epitope-specific T-Cell proliferation. **a**, Peptide incorporation onto the C-terminus of LF_N_ by sortase-mediated ligation. **b**, Illustration of translocation by protective antigen (PA) and the N-terminus of lethal factor (LF_N_). **c**, Biotinylated peptides **1**–**3**. **d**, Western blot analysis after PA-mediated translocation of LF_N_ **1**–**3** in CHO-K1 cells, with and without pre-treatment of cells with 20 µM lactacystin. (E–L) Flow cytometry analysis of Cell-Trace Violet (BV421)-labeled T cells (OT-1) after co-incubation with C57BL/6 dendritic cells (DCs), which were treated with PA (20 nM) and LF_N_-OVA **1**–**8** or **1G**–**8G** (1 µM, unless otherwise indicated). **e**, Representative histogram plots of T-cell proliferation from PA-translocation of LF_N_-OVA **3** and **6**. **f**–**g**, T-cell activation and proliferation of the individual responses from PA/LF_N_-OVA **1**–**8**, with and without lactacystin pre-incubation. **h**, Grouped pairwise comparison of the responses from panel **3g**. **i**, Pearson correlation of C-terminal degron index (CDI) vs. T cell proliferation (% Divided) from translocated LF_N_-OVA **1**–**8** (10, 1, and 0.1 µM). **j**, Grouped comparisons of the T-cell proliferation responses from (I). **k**–**l**, Pairwise comparisons of T cell proliferation after translocating: LF_N_-OVA **1**–**8** and **1G**–**8G**; and LF_N_-OVA **1**–**8** and **9**–**12**. Statistical analyses in **h**, **j**, **k**, and **l** are Welch’s t tests that were performed to compare T cell proliferation activity of the grouped responses, which indicated the comparisons were significantly different: P < 0.0332 (*); P < 0.0021 (**); P < 0.0002 (***); and P < 0.0001 (****). Data are representative of three independent experiments.

We measured the presence of translocated LF_N_ protein to evaluate whether proteasomal degradation is favored or disfavored based on CDI. For these studies, we combined LF_N_ with synthetic peptides **1**–**3** that show varying CDI values: 40, 71, and 80 (Fig. 3c). Proteasomal degradation was evaluated with CHO-K1 cells, which are an established cell line that express the anthrax receptors and enable PA-mediated binding and cytosolic protein translocation^37^. The peptides were evaluated with and without pre-treating CHO cells with the lactacystin proteasome inhibitor, followed by incubating the cells with PA (20 nM) and LF_N_ **1**–**3** (100 nM). Cytosolic extraction with a digitonin buffer, followed by Western blot analysis established the cytosolic fraction based on the presence of horizontal bands associated with ERK1/2, which are well-known MAP kinase proteins that are located in the cytosol (Fig. 3d)^41^. The cytosolic fraction was further established by the absence of a Rab5 band, which is a key protein that localizes in early endosomes^42^. Proteasomal degradation of LF_N_ **1**–**3** was established based on the intensity of horizontal bands associated with biotinylated protein, which become darker after lactacystin treatment. These studies establish that PA/LF_N_ successfully delivers peptides **1**–**3** into CHO-K1 cells, and that peptide degradation occurs in a proteasome-dependent fashion.

### Automated flow synthesis of vaccine epitopes

We used automated flow peptide synthesis (AFPS) to accelerate studies that relate proteasomal stability to epitope immunogenicity^43^. We synthesized antigen peptides derived from ovalbumin (OVA) that contain the OVA_257-264_ (SIINFEKL) epitope to impart immunogenicity^44,45^. We designed 20 peptide variants, which were prepared and conjugated to LF_N_ (Table 1). The peptides each comprise three N-terminal Gly residues for sortase-mediated ligation; native epitope residues from OVA_257–264_ (SIINFEKL) epitope for immunogenicity; and varying flanking residues for tuning degradation activity. A subset of eight peptides, called OVA **1**–**8**, contain native resides but only differ in length (13–28 amino acids); A subset of four peptides, called OVA **9**–**12**, contain mutated flanking residues that impart a high score (i.e., CDI > 50). Another subset of eight peptides, called OVA **1G**–**8G**, are homologues of OVA **1**–**8** but contain two additional Gly residues that impart a low score (i.e., CDI < 25)^27^. The peptides were prepared by the AFPS on HMPB-ChemMatrix resin, cleaved from the resin under acidic conditions, and purified by RP-HPLC^43^. Mass spectrometry (ESI) analysis showed the desired mass for the resulting LF_N_-OVA fusion proteins after sortase-mediated ligation.

**Table 1.**
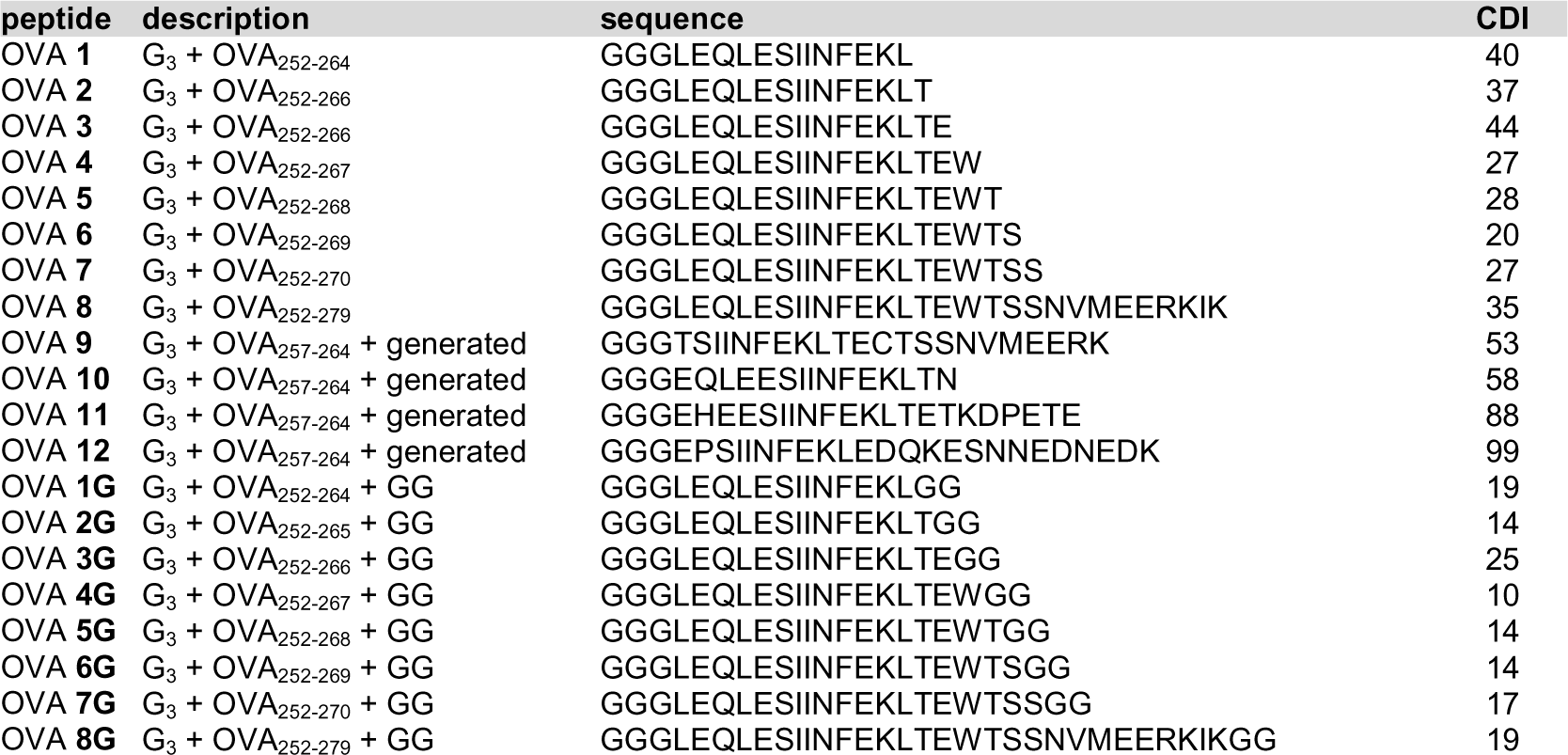
Summary of ovalbumin (OVA) peptides 1–12 and 1G–8G.

### Proteasomal stability reflects T cell epitope immunogenicity

We used the PA/LF_N_-OVA constructs to evaluate the influence of a degron sequence on immunogenicity. Primary dendritic cells (DCs) from mouse splenocytes (C57BL/6) were obtained, treated with lactacystin (20 µM) for 1 h, and co-incubated with PA/LF_N_-OVA constructs (Fig. 3e–k). Co-incubating the DCs with CellTrace Violet-labeled T cells (OT-1) followed by flow cytometry analysis revealed T-cell activation (24 h) based on upregulation of CD69 and CD137 markers, and also revealed T-cell proliferation (72 h) based on dilution of the dye (Fig. 3e).

Immunogenicity studies with the LF_N_-OVA epitopes established that T cell proliferation is dependent on proteasomal degradation (i.e., CDI). The proteasome dependance is reflected by diminished T cell activation in the presence of a proteasome inhibitor. The LF_N_-OVA epitopes were evaluated by pre-incubating murine DCs with and without 100 µM lactacystin, followed by evaluating proliferation of OVA-specific T cells (Fig. 3f,g). Grouping the activities from the LF_N_-OVA **1**–**8** constructs, with and without treatment of lactacystin, showed a significant decrease in immunogenicity due to the inhibited proteasome (Fig. 3h).

Titrating the LF_N_-OVA concentration revealed a dependence of concentration on proteasome-mediated immunogenicity. The concentration dependance is reflected by translocating LF_N_-OVA **1**–**8** epitopes at varying concentrations. At the lowest concentration (0.1 µM), LF_N_-OVA epitopes show a strong correlation (R^2^ = 0.85) between CDI and T cell proliferation (% divided). At higher concentrations (1 and 10 µM), the correlation steadily decreases (R^2^ = 0.68 and 0.04) due to saturation of the antigen presenting cells (Fig. 3i, j). These results suggest that degron activity is not only important for tuning epitope immunogenicity, but also for maximizing epitope efficacy at lower concentrations.

Mutating the flanking residues of the OVA epitope further reveals the influence of proteasomal degradation. LF_N_-OVA **1**–**8** show modest T cell proliferation activity that varies between each construct (Fig. 3k); LF_N_-OVA **1G**–**8G** show pronounced T cell proliferation that is nearly identical across all eight sequences (Fig. 3k); and LF_N_-OVA **9**–**12** show limited T cell proliferation (Fig. 3l). The mutated OVA sequences further demonstrate the influence of CDI on immunogenicity. Although the T cell epitope is identical across all sequences, the activity varies for both the native and mutated sequences. These variations appear to reflect the predicted CDI, in which the magnitude of T cell proliferation greatly depends on whether degradation is predicted to increase or decrease.

### Tuning proteasomal stability enhances immunogenicity

We further evaluated degron sequences through binary comparisons of high and low degradation activities. These comparisons shed light on epitope immunogenicity for other disease models (Fig 4). For each epitope, we generated randomized C-terminal sequences using a machine learning-based goal-search algorithm, in which the randomized sequences consist of 30,000-membered libraries. Each peptide comprises the native epitope sequences (8–9 amino acids) and N-terminal (5 amino acids) residues, but also contain randomized C-terminal residues (10 amino acids). Peptides were selected from the library that exhibit low (CDI_LO_) and high (CDI_HI_) degradation scores. LF_N_ conjugates were prepared by synthesizing the CDI_LO_ and CDI_HI_ peptides using automated flow peptide synthesis, followed by purification and conjugation to LF_N_.

**Fig. 4.**
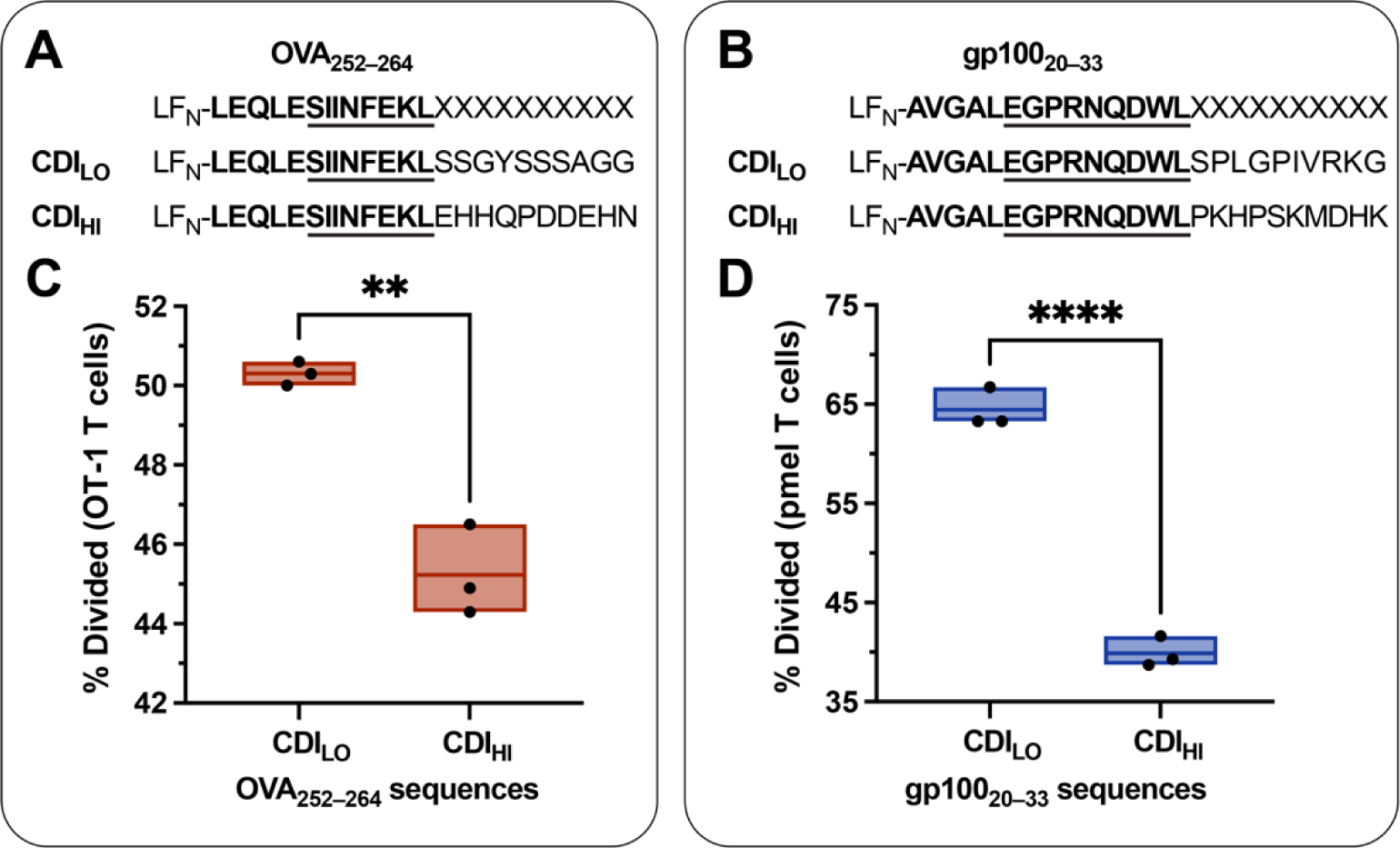
Promoting proteasomal degradation enhances epitope immunogenicity. Epitope sequences (8–9 amino acids) with varying proteasomal stabilities (CDI) adopted from: ovalbumin (OVA_252–264_) and glycoprotein 100 (gp100_20–33_). Each sequence comprised the native residues from the N-terminal (bolded) and epitope (underlined) regions, followed by randomized C-terminal residues (10 amino acids). The randomized residues (XXXXXXXXXX) were generated using a ML goal-search algorithm, which afforded diverse sequences (30,000-membered peptide library) with varying proteasomal stabilities, including CDI = <10 (CDI_LO_) and >60 (CDI_HI_). **a**, **b** Peptides from each group were selected and synthesized, each containing three N-terminal Gly residues (i.e., GGG-peptide) for conjugation to LF_N_ with SrtA*. **c**, **d**, Primary dendritic cells (C57BL/6) were treated with PA (20 nM) and the indicated LF_N_-CDI_LO_ or LF_N_-CDI_HI_ (1 µM), followed by co-incubation (72 h) with CTV-labeled T cells obtained from transgenic mouse models: OT-1 and human pre-melanosome protein (pmel). Statistical significance was calculated using one-way ANOVA to compare groups, which showed the indicated comparisons were significantly different: P < 0.0332 (*); P < 0.0021 (**); P < 0.0002 (***); and P < 0.0001 (****). Data are representative of at least three independent experiments.

Relative immunogenicity of the CDI_LO_ and CDI_HI_ peptides was evaluated with primary DCs (C57BL/6). The DCs were treated with PA and the LF_N_-CDI_LO_ and LFN-CDI_HI_ conjugates, followed by coincubation (72 h) with CellTrace Violet-labeled T cells obtained from transgenic mouse models: OT-1 and human pre-melanosome protein (pmel). Flow cytometry analysis of the T cells revealed CDI-dependent proliferation from the translocated epitopes into DCs: CDI_LO_ peptides show pronounced proliferation; CDI_HI_ epitopes show limited proliferation. This comparison shows that reducing the CDI can increase epitope-specific T cell proliferation through favoring proteasomal degradation.

## Discussion

Although proteasomal degradation is an established step in the antigen processing pathway, several limitations have precluded the development of epitope design rules thus far. The limitations include: incomplete characterization of the sequences that influence proteasomal degradation^46^; and insufficient cytosolic delivery into antigen presenting cells. As a result, the degradation propensity for many immunogenic epitopes remained unclear, including for established disease models associated with cancer, virus, and bacterial epitopes^31,47–49^.

Among clinically-studied vaccines, particularly personalized vaccines, only a small subset of the immunizing peptides demonstrated priming of cytotoxic T cells^2^. To evaluate whether proteasomal degradation influences T cell priming in clinical settings, we used the model to evaluate degron activity across three clinical trial studies of personalized vaccine peptides: two for melanoma^50,51^ and one for glioblastoma^52^. We evaluated the immunizing peptide sequences from these studies by dividing the results into two groups: presence (+) or absence (–) of CD8^+^ T cell responses after vaccination (Fig. 5). The two groups were then plotted against the CDI values (left Y axis), in which the shading of the individual points reported the HLA-binding score (right Y axis).

**Fig. 5.**
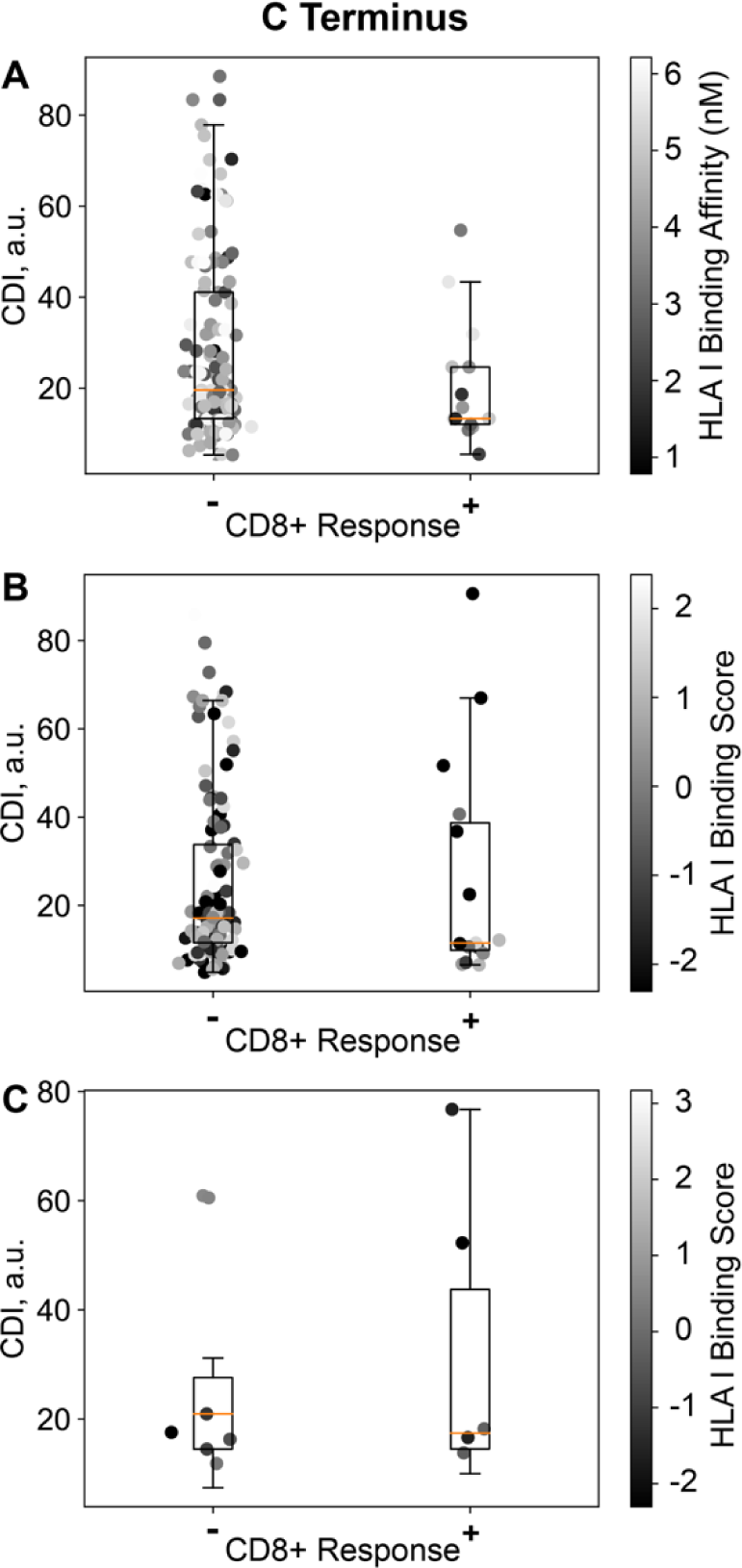
Retrospective analysis of clinically studied vaccine sequences. Vaccine sequences were evaluated for proteasomal degradation activity based on C-degron stability: **a** personalized peptide vaccines for melanoma; **b** personalized RNA vaccines for melanoma; and **c** personalized peptide vaccines for glioblastoma. CDI results are graphed using box plots (left Y axis); Individual data points are shaded based on reported binding affinity to HLA molecules (right Y axis). The prediction results were divided into two groups based on the successful (+) or unsuccessful (-) detection of a CD8+ T cell response after vaccination.

Several trends emerged from these retrospective studies. From Ott and co-workers (2017), sequences exhibiting low CDI showed CD8^+^ T cell activation, indicating that proteasomal degradation of peptide-based vaccines favors cytotoxic T cell responses (Fig. 5a); From Sahin and co-workers (2017), the sequences showed no difference between the unsuccessful sequences, which suggests that some RNA-encoded epitopes sequences can be degraded before completion of ribosomal synthesis (Fig. 5b); From Hilf and co-workers (2019), the successful sequences did not show a difference in mean CDI values, however, this studies comprised few immunizing sequences and limits the ability to draw conclusions (Fig. 5c). Taken together, these three studies suggest that degron activity is an important feature for designing vaccine epitopes. Nonetheless, further studies are needed to correlate the influence of vaccine formulation on the antigen degradation in vivo.

## Conclusion

This work provides a conceptual framework for combining degron sequences with vaccine epitopes. These sequences permit the design of epitopes to favor degradation or, alternatively, to favor stability. Although degron sequences are complex, machine learning accommodates these patterns for predicting degradation propensity. Incorporating degradation propensity into vaccine epitope design was shown to enhance epitope immunogenicity without altering the epitope. Degron sequences also enabled tuning of proteasomal degradation across disease models, particularly for designing the flanking residues of epitope sequences associated with model antigens, tumor antigens, and personalized neoantigens.

Essential to this work was the use of the PA/LF_N_ delivery system. PA/LF_N_ facilitated translocation of epitope sequences into cells for evaluating degradation propensity. Further analysis showed that relative proteasomal stability correlates with immune activation activity. These studies show promise for future efforts to improve vaccine epitope designs against tumors and pathogens in whole animal models and clinical settings.

## Materials and Methods

Fmoc-protected L-amino acids used for peptide synthesis were purchased from Novabiochem. Peptide synthesis couplings were performed with 1-[bis-(dimethylamino)methylene]-1H-1,2,3-triazolo[4,5-b]-pyridinium 3-oxid hexafluorophosphate (HATU) and (7-Azabenzotriazol-1-yloxy)tripyrrolidinophosphonium hexafluorophosphate (PyAOP), which were purchased from P3 Biosystems. Dimethylformamide, piperidine, diisopropylethylamine, trifluoroacetic acid, and triisopropylsilane were purchased from VWR or Millipore Sigma. Antibodies for flow cytometry were purchased from BioLegend. Media for cell culture were purchased from ThermoFisher Scientific. Tissue culture was performed with RPMI 1640 Medium, GlutaMAX™ Supplement, Fetal Bovine Serum, qualified, One Shot™ format. Penicillin-Streptomycin (10,000 U/mL) was purchased from ThermoFisher Scientific. Western blots were performed with nitrocellulose membranes (GE), filters (Bio-Rad Laboratories, Inc.), and PBS blocking buffer (LI-COR Biosciences). Primary and secondary antibodies for visualization of the bands include: Erk1/2 (Cell Signaling), goat anti-mouse IRdye680 (LI-COR Biosciences), and streptavidin IRdye680 (LI-COR Biosciences).

### General Equation

The following equation uses: a linear combination of bin populations (e.g., bin1, bin2, bin3, and bin4), which was obtained from Elledge and coworkers; exponentially increasing coefficients (e.g., 0, 1, 10, and 100), which reflect the exponential scale of the original data (i.e., flow cytometry). The resulting equation, which we call the C-terminal Degron Index (CDI), relates proteasomal degradation activity to a numerical score that ranges from 0 to 100.

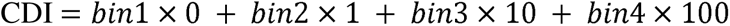

### Synthesis and purification of antigen peptides

Peptides were synthesized on a 0.1 mmol scale by automated flow peptide synthesis. Peptide synthesis was performed on ChemMatrix resin with a 4-(4-Hydroxymethyl-3-methoxyphenoxy)butyric acid (HMPB) linker (200 mg, 0.5 mmol/g, 100–200 mesh). The first amino acid (1 mmol, 10 equiv.) was manually coupled to the resin with DIC (0.5 mmol, 78 µL) and DMAP (0.01 mmol, 50 µL of a 0.2M solution in DMF) in 3.17 mL of DMF. The resin suspension was incubated overnight (16–24 h), then was drained and rinsed three times with DMF (5 mL). Subsequent amino acids were added by automated flow peptide synthesis. After the syntheses were complete, peptide cleavage and global deprotection was performed with a solution of trifluoroacetic acid, water, ethane dithiol, and triisopropyl silane (94/2.5/2.5/1). Purification was achieved by preparative RP-HPLC with Agilent Zorbax SB-C18 Prep HT column (21.2 mm × 250 mm, 7 μm) at a flow rate of 15 mL/min using a gradient with water and acetonitrile containing 0.1% TFA. Pure HPLC fractions were pooled and lyophilized. The purified peptides were analyzed as 0.01 mg/mL solutions (50:50 CH_3_CN in H_2_0 with 1% formic acid) by LC/MS on an Agilent 6550 ESI-Q-TOF mass spectrometer equipped with an Agilent Zorbax 300SB-C3 column (2.1 mm × 150 mm, 5 µM) with a 1–91% gradient of CH_3_CN in H_2_O with 0.1% formic acid and a flow rate of 0.5 mL/min.

### Protein expression and purification

#### Protective antigen (PA)

This protein was expressed in *B. anthracis* strain BH500 from a pYS5 plasmid, which gave PA in high yields with limited endotoxin. Cultures containing plasmid were grown in FA medium containing 10 μg/ml of kanamycin at 37°C for 14 h. The cultures were cooled and supplemented with 2 μg/ml of AEBSF [4-(2-Aminoethyl)-benzenesulfonylfluoride HCl], and then centrifuged at 4550 g for 30 min. All subsequent steps were performed at 4°C. The supernatants were filter sterilized and supplemented with 5 mM EDTA. Solid ammonium sulfate was added to the supernatants to obtain 40% saturation. Phenyl-Sepharose Fast Flow (low sub) (GE Healthcare Life Sciences, Uppsala, Sweden) was added and supernatants gently mixed in 4°C for 1.5 h. The resin was collected on a fritted-disk funnel and washed with buffer containing 1.5 M ammonium sulfate, 10 mM Tris HCl, and 1 mM EDTA (pH 8.0). Protein was eluted with 0.3 M ammonium sulfate, 10 mM Tris HCl, and 1 mM EDTA (pH 8.0), precipitated by adding an additional 30 g ammonium sulfate per 100 mL eluate, and centrifuged at 18,370 g for 20 min. Protein was resuspended in 5 mM HEPES, 0.5 mM EDTA (pH 7.5), followed by loading onto a Q-Sepharose Fast Flow column (GE Healthcare Life Sciences). Protein was eluted with a 0–0.5 M NaCl gradient in 20 mM Tris–HCl, 0.5 mM EDTA (pH 8.0). Protein-containing fractions were identified by SDS-PAGE at 165 V for 36 min on an Invitrogen Bolt™ 4–12% Bis-Tris Plus Gel with Bolt™ MES SDS Running Buffer (1x). Gels were visualized by SimplyBlue™ SafeStain (Coomassie). The clean fractions were pooled and buffer-exchanged into 10 mM Tris, pH 7.5, 150 mM NaCl, and 0.5 mM EDTA. Protein was concentrated as necessary, flash-frozen in liquid nitrogen, and stored at –80 °C. The exact mass of the purified protein was confirmed by LC/MS on an Agilent 6550 ESI-Q-TOF mass spectrometer equipped with an Agilent Zorbax 300SB-C3 column (2.1 mm × 150 mm, 5 µM) with a 1–91% gradient of CH_3_CN in H_2_O with 0.1% formic acid and a flow rate of 0.5 mL/min.

#### N-terminus of Lethal Factor (LF_N_)

This protein was expressed in BL21(DE3) *E. coli*, which was purchased from New England Biolabs. The protein was expressed at New England Regional Center of Excellence/Biodefense and Emerging Infectious Diseases (NERCE) and was purified. LF_N_ was expressed as the SUMO-LF_N_-LPSTGG-H_6_ construct in a Champion pET-SUMO vector. LF_N_ was isolated from *E. coli* pellets by suspension in Tris buffer (20 mM Tris, 150 mM NaCl, pH 8.5), lysis by sonication, and purification with a HisTrap FF Ni-NTA column. Purified fractions of PA and LF_N_ were analyzed by SDS-PAGE at 165 V for 36 min on an Invitrogen Bolt™ 4–12% Bis-Tris Plus Gel with Bolt™ MES SDS Running Buffer (1x). Gels were visualized by SimplyBlue™ SafeStain (Coomassie). Clean fractions were pooled and concentrated with Amicon® Ultra-15 Centrifugal Filter Units.

### Sortase-mediated ligation

Semi-synthetic LF_N_ protein constructs were prepared by enzymatic ligation using SrtA*. LF_N_-LPSTGG-H_6_ (80 μM) was combined with synthetic peptides (800 μM) in PBS buffer (pH 8.5, endotoxin free, Corning), in addition to adding a 50x dilution of 0.5 M CaCl_2_ reaction buffer (freshly prepared, endotoxin free) to give a final concentration of 10 mM CaCl_2_. The reaction mixture was gently rotated for 45 min., followed by adding triple-rinsed Ni-NTA agarose beads (50 µL per mg of protein), which enabled isolation of the enzyme: H_6_-SrtA*; reacted starting material: GG-H_6_; and unreacted starting material: LF_N_-LPSTGG-H_6_. Collection of the reaction supernatant was achieved by centrifugation of the reaction mixture (30 sec × 16,000 rpm), followed by successive rounds of rinsing with PBS (3 × 0.5 mL). To the combined rinses was added aqueous EDTA (0.5 M, pH 7.5, 100 µL, freshly prepared, endotoxin free) to sequester the CaCl_2_. The mixture was pushed through a 0.2 µm syringe filter, then buffer exchanged using Amicon® Ultra-15 Centrifugal Filter Units (MSCO = 30 kDa) to remove the excess peptide. The purified peptide-conjugated LF_N_ was analyzed by SDS-PAGE at 165 V for 36 min on an Invitrogen Bolt™ 4–12% Bis-Tris Plus Gel with Bolt™ MES SDS Running Buffer (1x). Gels were visualized by SimplyBlue™ SafeStain (Coomassie). The exact mass of the purified protein was confirmed by LC/MS on an Agilent 6550 ESI-Q-TOF mass spectrometer equipped with an Agilent Zorbax 300SB-C3 column (2.1 mm × 150 mm, 5 µM) with a 1–91% gradient of CH_3_CN in H_2_O with 0.1% formic acid and a flow rate of 0.5 mL/min.

### LC-MS Protein Characterization

Protein (50 ng) was loaded onto an Agilent Zorbax 5 μm 300SB-C3 column (2.1 × 150 mm) and was eluted with a gradient of 1–91% CH3CN in H2O with 0.1% FA and a flow rate of 0.5 mL/min. The protein was detected on an Agilent 6550 ESI-Q-TOF mass spectrometer.

### Western blot

Proteasome-mediated degradation was examined by Western Blot analysis. These experiments were performed by plating CHO-K1 cells at 2 × 10^5^ cells/well in a 12-well tissue culture plate. After incubating for 16 h at 37 °C and 5% CO_2_, the cells were resuspended in media treated with or without 50 µM lactacystin. After incubating for 1 h, the cells were resuspended in media containing PA (50 nM) and an LF_N_ construct (LF_N_ **1**–**3**). After incubating for 24 h, the wells were washed with PBS and trypsonized (0.25% trypsin–EDTA) for 5 min at 37 °C and 5% CO_2_. The cell pellets were treated with lysis buffer (50 µg/mL digitonin, 75 mM NaCl, 1 mM NaH_2_PO_4_, 8 mM Na_2_HPO_4_, 250 mM sucrose, and Roche cOmplete^TM^ protease inhibitor cocktail, pH 7.7) for 10 min on ice, then pelleted for 5 min at 4 °C and 16,000 rcf. The lysates were filtered using AcroPrep Filter Plates with Bio-Inert Membranes (Pall Life Sciences), then separated by SDS-PAGE and transferred onto Western Blot membranes. The membranes were developed using optimized conditions from prior expeirments (not shown). Membrane blocking was achieved by incubating for 1 h in PBS containing 5% LI-COR blocking buffer. Primary antibodies were incubated for 2 h in a solution of TBST (0.05%) and LI-COR blocking buffer (5%) containing the following antibodies: anti-ERK1/2 (Cell Signaling Technologies; dilution factor: 1/2000) and streptavidin IRdye680 (LI-COR Biosciences; dilution factor: 1/2000). The goat anti-rabbit IRdye680 secondary antibody (LI-COR Biosciences; dilution factor 1/5000) was incubated for 1 h in TBST (0.05%) and LI-COR blocking buffer (5%). The membrane was imaged on an Odyssey Imaging System (LI-COR Biosciences).

### Endotoxin testing and removal

Protein endotoxin levels were measured at 0.045 µg/mL using single-use cartridges (0.05 EU/mL, PTS2005) and the Endosafe® nexgen-PTS™ reader (Charles river). If endotoxin levels were > 0.015 EU/µg, established procedures to remove endotoxin were followed. In particular, Pierce™ High-Capacity Endotoxin Removal Resin was used to reduce endotoxin levels for peptide-conjugated LF_N_ constructs. Prior to use, a 1-mL aliquot of resin was centrifuged (14000 rpm × 5 min) and resuspended in an equivalent volume of endotoxin-free phosphate-buffered saline (PBS). The resuspended resin was then added to each protein sample at a 1:4 or 1:8 v/v ratio (resin volume/protein volume). After mixing gently for 1 hr at room temperature, the resin was removed by filtering the samples through a 0.2 μm filter. This procedure provided high protein recovery (∼90%) and was repeated (1–3x) until sufficient endotoxin levels were reached (≤ 0.015 EU/µg).

### Murine immune response

Animal studies were carried out under an institute-approved IACUC protocol following federal, state, and local guidelines for the care and use of animals. C57BL6/J and OT-1 mice were procured from The Jackson Laboratory. Six-to 12-week-old female mice were used for these studies. Antibodies for flow cytometry studies were purchased from Biolegend: CD69 (clone H1.2F3); CD137 (clone 17B5); CD3 (clone 17A2); CD90.1 (clone OX-7); and CD8a (clone 53-6.7). Experiments with murine splenocytes utilized freshly harvested spleens from naïve mice. Dendritic cells were isolated with the EasySep^TM^ Mouse Pan-DC Enrichment Kit II (Stem Cell Technologies). T Cells were isolated with the EasySep^TM^ Mouse T Cell Isolation Kit (Stem Cell Technologies). Antibody staining was performed at a dilution of 1:100 for 25 minutes at 4 °C in the presence of mouse Fc block (TruStain FcX™, anti-mouse CD16/32, BioLegend) in PBS containing 5% FBS. Cell proliferation was monitored using CellTrace™ Violet Cell Proliferation Kit (ThermoFisher Scientific). Viability was assessed by LIVE/DEAD Fixable Aqua (Life Technologies). Cells were analyzed using BD LSR Fortessa. Data were analyzed using FlowJo v10. Peptides for restimulation were used at a concentration of 10 mg/mL with sequences as follows: OVA_257–264_, SIINFEKL and gp100_20–33_, EGPRNQDWL.

### Mixed lymphocyte reaction

Murine splenocytes were freshly isolated from mouse spleens by homogenization over a mesh filter (100 µm) in RPMI 1640 Medium (GlutaMAX™ Supplement, 10% Fetal Bovine Serum, and 1% Penicillin-Streptomycin). Single-cell suspensions were created by re-filtering the cells through a cell strainer (45 µm). For C57Bl/6 mice, dendritic cells (DCs) were enriched by magnetic isolation using the EasySep^TM^ Mouse Pan-DC Enrichment Kit (Stem Cell Technologies). DCs were plated into 96-well U-bottom plates (100 × 10^6^ cells/well), followed by incubation with or without lactacystin (20–100 µM), at 37 °C and 5% CO_2_. After 4 h, the DCs were centrifuged (5 min. × 400 rcf) and resuspended in the protein treatments. The DCs were incubated with the proteins for 1 h, followed by centrifugation and resuspension in T cells (50 × 10^6^ cells/well). During the DC incubations, T cells from the transgenic mouse splenocytes were enriched by magnetic isolation using the EasySep^TM^ Mouse T Cell Isolation Kit (Stem Cell Technologies). T cells were centrifuged and resuspended in PBS, followed by treatment with CellTrace™ Violet Cell Proliferation Kit (ThermoFisher Scientific) according to the manufacturer’s protocol. T cells were centrifuged and resuspended in RPMI media, plating with the DCs. After 24 h, upregulation of early-activation markers (CD69 and CD137) were evaluated. After 72 h, the magnitude of the T cell proliferation was evaluated by flow cytometry.

### Statistical analysis

Results from mixed lymphocyte reactions were analyzed using GraphPad Prism software. Data from protein constructs were grouped within the same experiment, then analyzed by unpaired t test with Welch’s correction (assumes that standard deviations are not equal).

### Representation of peptides

The peptides were represented as a matrix of extended connectivity fingerprints of individual amino acids, generated using RDKit (http://www.rdkit.org/). For the individual fingerprints, we used a radius of 3, and 128 bit-size. For the peptides, we considered only the last 24 amino acids, and used ‘left’ padding for shorter sequences.

### Machine learning

We developed a multi-layer Conv1D model using TensorFlow, and optimized the hyperparameters over 1000 iterations using SigOpt (https://sigopt.com/). The hyperparameter optimization minimized the RMSE as the objective function. After the hyperparameter optimization, we used the hyperparameters for the top 5 models, and re-trained 5 models each with different random seeds. Finally, we used 25 models (5 top-hyperparameter x 5 random-seed) for the prediction, analysis and screening of different peptides.

### Genetic algorithm for generation of new peptides

We used directed evolution to generate new peptides. For peptides of different lengths, we used a random seed of the desired length from the training dataset as the C-terminus degron, keeping the N-terminus and epitope constant. Then, we performed randomized single and multi-site swapping of amino acids. In the single-site swapping, we randomly selected a particular position at the C-terminus and replaced it with a random residue. Similarly, in the multi-site swapping, we selected a string of amino acids and replaced it with another random string from the training dataset with the same length. The objective of the genetic algorithm was to increase the CDI score.

### Selection of peptides for experimental evaluation

As a part of the genetic algorithm-based optimization, we obtained new peptides having a wide range of predicted CDI scores. We also calculated the Grand Average of Hydropathy (GRAVY) score and iso-electronic points (pI) to infer ease-of-experimental synthesis and purification. The new peptides were filtered with a GRAVY score of <= 0.1, and with a pI either between 4 and 6.5 or between 7.5 and 10.

## Acknowledgements

This work was funded by the 2018 Bridge Project between the Koch Institute and Dana-Farber/Harvard Cancer Center to B.L.P. and C.J.W. This work was also supported by funding (to N.L.T.) by the Koch Institute Ludwig Foundation, National Cancer Institute (F32-CA239362), and National Institute of General Medical Sciences (5-P20-GM109091). D.J.I. is an investigator at the Howard Hughes Medical Institute. D.B.K. is supported by 1R01HL157174-01A1. This work was supported in part by the NERCE facility (Grant: U54-AI057159) for expression of toxin proteins and by the Koch Institute Cancer Center Support (Core) Grant P30-CA14051 from the National Cancer Institute. The authors acknowledge the Koch Institute’s Robert A. Swanson (1969) Biotechnology Center for technical support, specifically the Preclinical Modeling and Flow Cytometry facilities. The authors also thank M. Bakalar, A. Callahan, E. Fritsch, N. Hacohen, A. Rondon, and A. Loas, for providing thoughtful feedback throughout the development of the experiments and the manuscript.

## Author contributions

N.L.T., S.M., D.B.K., C.J.W., R.G.B., B.L.P. designed the research. N.L.T., S.M., M.M., J.R., N.L., W.A., D.S., and F.T. performed all experiments. N.L.T. and S.M. analyzed the data and wrote the manuscript. All authors discussed the manuscript.

## Data/Code Availability

We have made the data and code available at https://zenodo.org/record/8338338.

